# Optimizing calcein marking in the common periwinkle (*Littorina littorea*) for repeated-measures and mark-recapture studies

**DOI:** 10.64898/2026.04.10.717815

**Authors:** Greta K. Ahlefeld, Catherine E. Benavides, Margaret A. Chioffi, Finnian Furtney, Stephanie Goerck de Carvalho Macedo, Christina E. H. Korn, Gabe Marra-Perrault, Eamon A. F. McGlashan, Lily A. Watts, Khalil J. Wilkinson, Christopher D. Wells

## Abstract

Calcein is a fluorescent marker commonly used to label growing calcified structures in marine organisms, but its efficacy is species- and context-specific. We evaluated calcein marking success and survival in the common periwinkle (*Littorina littorea*) during winter in the Gulf of Maine. Snails were immersed for 24 h in seawater containing 0, 50, or 100 mg L^-1^ calcein and scored for fluorescent marks 22 days later. Overall marking success was low (12.5% of exposed snails evaluated) but was strongly size-dependent: each 1 mm increase in shell length reduced the odds of acquiring a mark by 27%. Among exposed snails, higher calcein concentration (100 mg L^-1^) produced significantly brighter marks than the lower concentration (50 mg L^-1^). Survival was 100% across all treatments. The low overall marking rate likely reflects suppressed shell growth at winter temperatures. We recommend 100 mg L^-1^ calcein with a 24-h immersion for marking *L. littorea* and suggest that marking during warmer months would improve efficacy across a broader size range.

## 1. Introduction

Keeping track of individuals over time is a persistent challenge in marine studies. Reliable individual marking underpins estimates of growth and survival, tests of carryover effects, and any experimental design that requires repeated measurements on the same organism. Although a variety of marking techniques exist for shelled invertebrates, each carries practical and biological limitations. External paints and tags can abrade or be lost as shells wear, impede movement, become unreadable after extended periods, or increase susceptibility to visual predators (Henry and Jarne, 2007; Levin, 1990; MacGregor et al., 2023). Incorporated into hard tissues, fluorescent labels avoid many of these limitations by creating an internal timestamp that persists as the organism grows, enabling unambiguous assessment of new shell growth and supporting non-lethal mark-recapture and repeated-measures designs.

One such fluorescent label is calcein, a water-soluble fluorochrome that binds to calcium during mineralization and is excited by blue light. When viewed through a yellow longpass transmission filter, structures with calcein appear bright green. It has been used widely in fishes, bivalves, and gastropods, but optimal concentrations and exposure durations are strongly species- and context-specific, reflecting tradeoffs among mark intensity, retention, and animal welfare (e.g., Moran, 2000; van der Geest et al., 2011; Wilson et al., 1987). As a result, concentration-response relationships cannot be assumed without species-level evaluation of both efficacy and potential adverse effects (e.g., Wells and Sebens, 2017).

In this study, we evaluated survival and staining efficacy of calcein on the common periwinkle, *Littorina littorea*. This snail is one of the most conspicuous and important grazers on temperate North Atlantic rocky shores. *L. littorea* can change algal diversity and abundance, shift ecosystems from soft-bottom to hard-bottom habitats, and alter the behavior of other intertidal snails (Bertness, 1984; Brenchley and Carlton, 1983; Lubchenco, 1978). Its abundance, ease of collection, and straightforward husbandry have made it one of the most experimentally tractable intertidal gastropods, with extensive use in grazing ecology, population biology, invasion biology, and studies of shell growth and phenotypic plasticity (e.g., Behrens Yamada and Mansour, 1987; Blakeslee and Byers, 2008; Kemp and Bertness, 1984; Lubchenco, 1978; Wells et al., 2023). A dependable, unambiguous, and non-lethal marking protocol would enable short-term growth assays, mark-recapture experiments, and lab-to-field manipulations. Yet species-specific calcein performance, particularly survival during exposure and the clarity of the resulting shell mark, has not been empirically tested for *L. littorea*. Our study provides the concentration-response data needed to support future growth and demographic experiments with *L. littorea* and establishes a usable protocol while clarifying tradeoffs between mark visibility and potential adverse effects.

## 2. Methods

*Littorina littorea* (N = 2,004) were hand-collected on December 8, 2025, from the rocky intertidal zone (+0.5 to +1.5 m relative to mean low water) at the Bowdoin College Schiller Coastal Studies Center, Harpswell, Maine (43.7918 N, 69.9576 W). All snails within an 8 x 8 m area were collected to obtain a broad size distribution representative of the local population. Length from spire apex to the farthest point of the aperture, herein referred to as shell length, ranged from 4.1 to 38.03 mm. Snails were haphazardly allocated to twelve 5-gallon buckets (167 snails per bucket), each containing 10 L of ambient, 150-µm filtered seawater. Buckets were continuously aerated and immersed in 4 cm of flowing seawater in a sea table to maintain ambient temperature within the containers.

### 2.2 Calcein exposure

Concentrated calcein (CAS No. 154071-48-4) stock solutions were prepared by dissolving the calcein in freshwater at concentrations of 50 g L^-1^ and 100 g L^-1^. Buckets were randomly assigned to one of three treatments (N = 4 per treatment): control, low concentration (50 mg L^-1^), or high concentration (100 mg L^-1^). To achieve the target concentrations, 10 mL of the appropriate stock solution was added to each treatment bucket; control buckets received 10 mL of freshwater to equalize any effect of freshwater addition. Snails were immersed for 24 h without food.

After the 24-h immersion, snails were transferred in their original treatment groups to square plastic mesh basket cages (20 x 20 cm at the top, 14 x 14 cm at the bottom). A second inverted basket was secured to the top of each cage to prevent escape. The 12 cages were placed in a sea table with continuously flowing ambient seawater, partially submerging the cages. Snails were fed sugar kelp (*Saccharina latissima*) *ad libitum* for the remainder of the experiment. Water temperature during the experiment ranged from -1.3 to 2.9°C (average 1.0°C).

### 2.3 Assessment of fluorescent marks

Twenty-two days after immersion, shells were examined externally for fluorescent marks. Marks were visualized using an ULTRAFIRE H-B3 blue LED flashlight (470 nm peak excitation) viewed through DEX FIT SG210 yellow-lens safety glasses, which filter out most of the blue light, allowing clearer assessment of calcein fluorescence. Each snail was assigned to one of three ordinal categories: no mark (no visible fluorescence), light mark (any visible fluorescent mark, Fig. 1B), or bright mark (a continuous fluorescent band exceeding 1 mm in length, Fig. 1D). A single observer (CDW) conducted all scoring. Shell length was measured with digital calipers to the nearest 0.01 mm. To evaluate whether shell length influenced marking success, we used a stratified scoring protocol: the 10 smallest snails per bucket were scored first, followed by 10 randomly selected individuals from the remaining snails, yielding 20 scored snails per bucket (N = 240 total). This design ensured adequate representation of small, potentially faster-growing individuals that might otherwise be underrepresented in a purely random sample. Representative specimens were photographed with a Leica DFC450 C camera mounted on a Leica M165 FC fluorescent stereo microscope illuminated with a NightSea Royal Blue Light and Filter Set to photograph calcein fluorescence.

**Figure 1.**
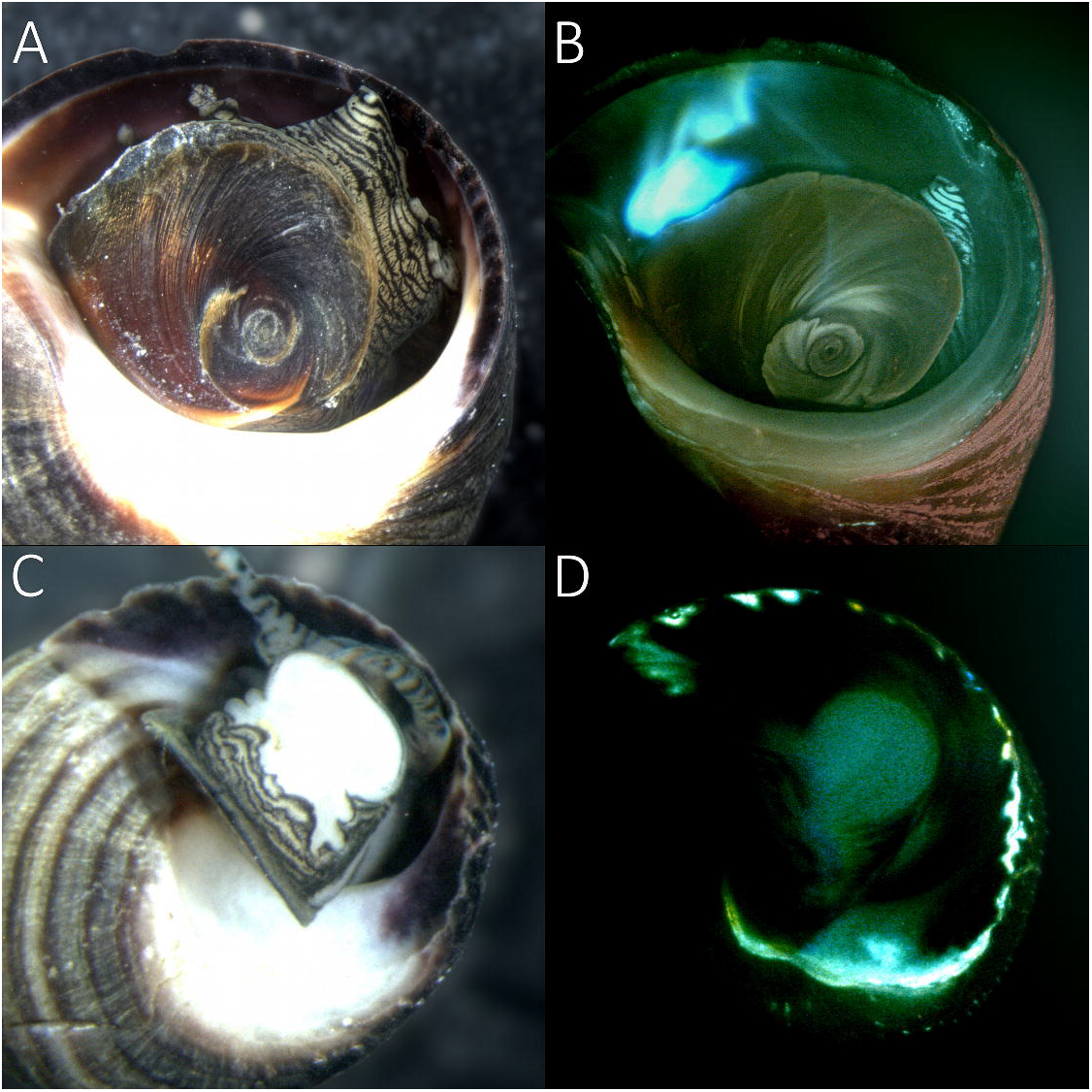
Representative *Littorina littorea* photographed under natural light (A and C) and blue light (470 nm) with a yellow longpass filter (B and D). The large snail (25.30 mm shell length, A and B) was scored as unmarked under the field protocol but shows faint calcein fluorescence (light blue) at the growing edge, in the body tissue, and in the fluid from the mantle cavity under a dissecting microscope. The small snail (5.72 mm shell length, C and D) shows a bright calcein mark along the shell margin.

### 2.4 Statistical Analyses

All statistical analyses were conducted in R v4.5.2 (R Core Team, 2025). Because our scoring protocol specifically targeted the smallest snails in addition to randomly selected individuals, we included shell length as a continuous covariate in all models to control for size-dependent marking success and to account for the stratified sampling design.

To test whether treatment exposure (none vs. exposed) and snail size affected the probability of acquiring any mark, we used generalized estimating equations (GEE) with a binomial error distribution and logit link (R package geepack 1.3.13, Højsgaard et al., 2005). We chose a GEE framework over a generalized linear mixed model (GLMM) because the number of clusters (N = 12 containers) was small. While GLMMs estimate cluster-specific random effects, their variance components can be difficult to estimate reliably with few clusters. GEEs focus on the marginal (population-averaged) effect of treatment and, when combined with small-sample bias corrections, provide robust inference for designs with as few as 12 clusters (Liang and Zeger, 1986; Mancl and DeRouen, 2001). We specified an independence working correlation structure. To ensure valid inference with a limited number of clusters, we assessed statistical significance using Wald tests with CR3 bias-corrected robust standard errors and Satterthwaite degrees of freedom (R package clubSandwich 0.6.2, Pustejovsky, 2026). 95% confidence intervals (CIs) for the predicted probabilities were calculated analytically using the CR3 robust covariance matrix.

To test whether treatment concentration (50 or 100 mg L^-1^) and snail size affected the intensity of the mark among exposed snails, we used an ordinal logistic regression with a logit link to account for the ordered nature of the response variable (R package ordinal 2023.12-4.1, Christensen, 2023). We restricted this analysis to snails exposed to calcein only. Consistent with the binary analysis, we accounted for clustering within containers using robust variance estimation rather than random effects to ensure stable estimation with a small number of clusters. We assessed statistical significance using cluster-robust standard errors (type HC1, R package sandwich 3.1-1, Zeileis, 2004; Zeileis et al., 2020). 95% CIs were generated using a cluster-based bootstrap with 1,000 iterations to preserve the correlation structure within containers.

## 3. Results

Treatment exposure was the primary determinant of marking success (β = -43.59, *SE* = 0.83, *t*(3.0) = -52.4, *p* < 0.001; Fig. 2). No snails scored from the control group acquired a mark (0%; 0/80), compared to 12.5% (20/160) of snails from the groups exposed to calcein. Snail size was also a significant predictor of marking success; larger snails were significantly less likely to acquire a mark (β = -0.31, *SE* = 0.05, *t*(6.8) = -6.84, *p* < 0.001; Figs. 1 and 2). For every 1 mm increase in shell length, the odds of acquiring a mark decreased by approximately 27% (*OR* = 0.73).

**Figure 2.**
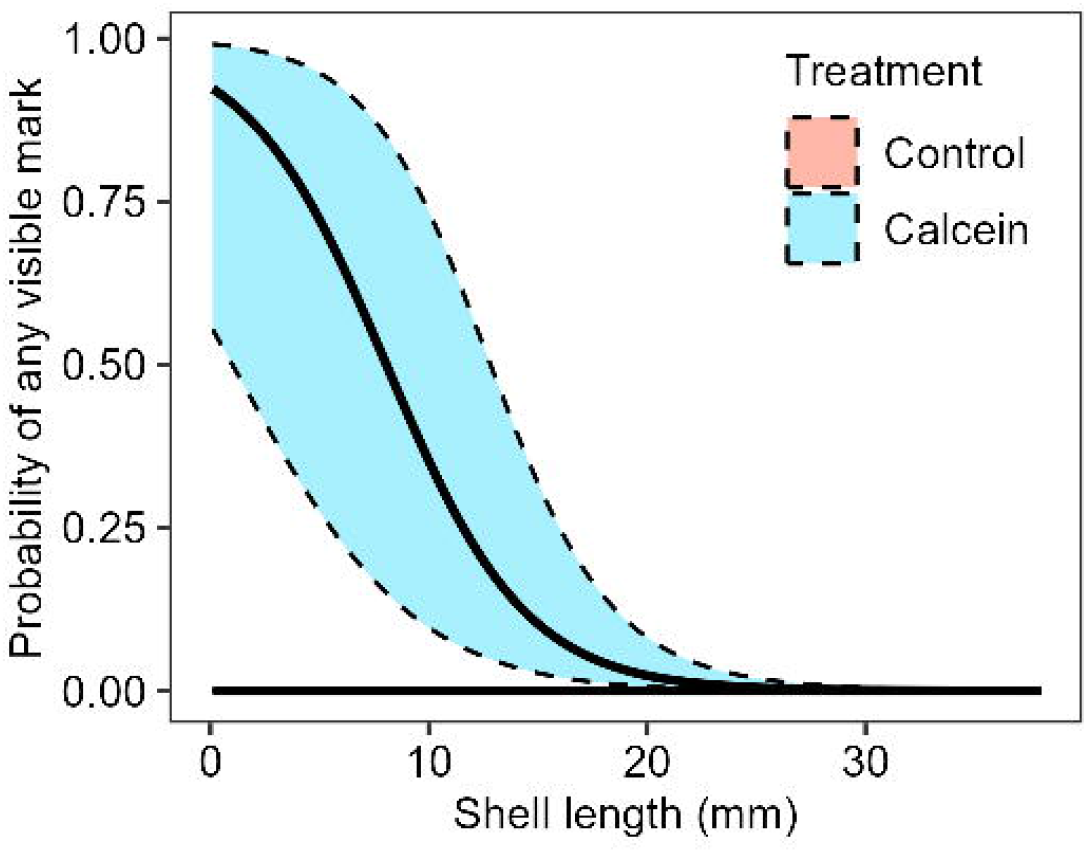
The effect of *Littorina littorea* shell length and exposure to calcein on the probability of observing a mark. Bold lines are fitted predictions from the generalized estimating equations with 95% confidence intervals represented by dashed lines and shading.

Among the 160 snails sampled from the calcein treatment groups, mark intensity was significantly influenced by calcein concentration and snail size. Snails in the high concentration treatment exhibited significantly brighter marks than those in the low concentration treatment (100 vs. 50 mg L^-1^; β = 0.92, *SE* = 0.39, *z* = 2.36, *p* = 0.020; Fig. 3). Shell length negatively affected mark intensity, with larger snails receiving significantly lighter marks (β = -0.37, *SE* = 0.08, *z* = -4.79, *p* < 0.001). For every 1 mm increase in length, the odds of a snail being in a brighter mark category decreased by approximately 31% (*OR* = 0.69).

**Figure 3.**
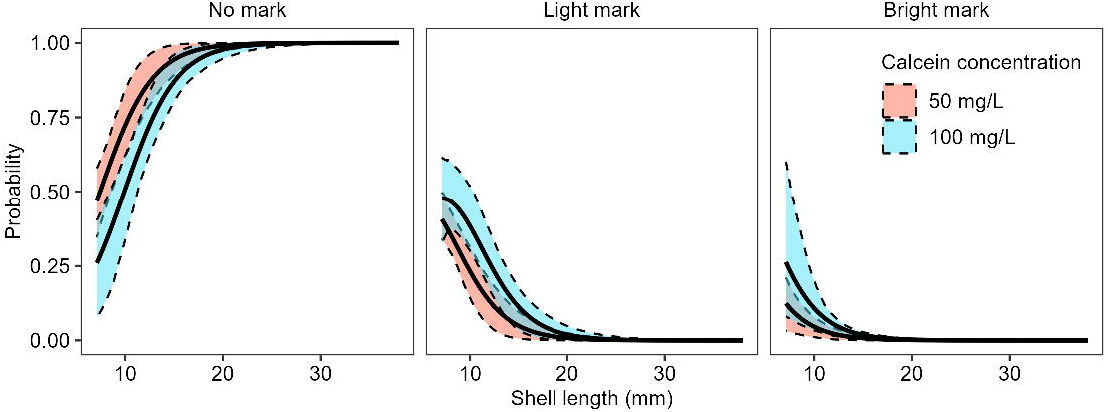
The effect of *Littorina littorea* shell length and calcein concentration on the probability of observing varying qualities of a mark. Bold lines are fitted predictions from the ordinal logistic regression with 95% confidence intervals represented by dashed lines and shading.

Exposure to calcein did not affect survival or behavior. All snails survived throughout the 22-day experiment. Snails remained active during the 24-h immersion and readily consumed kelp (*S. latissima*) during the recovery period. Additionally, fluorescent marks on exposed snails persisted throughout the recovery period. Calcein fluorescence was also visible within the soft tissue of marked individuals, though we did not quantify this systematically. Examination under a dissecting microscope revealed faint fluorescence in snails scored as unmarked under the field protocol (Fig. 1B), suggesting that the handheld method provides a conservative estimate of marking success.

## 4. Discussion

Calcein produced visible fluorescent marks in *L. littorea* following a 24-h immersion, but overall marking success was low (12.5% of exposed snails evaluated), with both mark acquisition and mark intensity strongly dependent on snail size and calcein concentration. These results are consistent with the expectation that calcein incorporation requires active shell deposition and therefore depends on the rate at which new shell material is being laid down during the exposure window.

The pronounced size effect, whereby each 1 mm increase in shell length reduced the odds of acquiring any mark by approximately 27% and of receiving a brighter mark by approximately 31%, likely reflects allometric growth patterns. Smaller *L. littorea* deposit new shell material at higher relative rates than larger conspecifics (Moore, 1937), creating more opportunity for calcein to be incorporated into the growing edge during a fixed exposure period. This pattern is consistent with findings in other calcein-marked mollusks: Moran (2000) found that newly hatched *Nucella ostrina* retained bright calcein marks after short immersions and van der Geest et al. (2011) reported variable marking success in adult bivalves that they attributed to individual differences in growth rate. The low overall marking rate in our study likely reflects cold temperatures (-1.3 to 2.9°C) suppressing growth rates across all size classes, limiting calcein incorporation even in the smaller, faster-growing individuals that would otherwise be the most likely to mark. Other intertidal littorinid snails display seasonal cold acclimation, reducing overall metabolic capacity and enzymatic activity (Sokolova and Pörtner, 2001). Coupled with strong seasonal shell growth in *L. littorea* (Moore, 1937), marking during winter represents a conservative test of calcein performance.

The significant effect of concentration on mark intensity, with 100 mg L^-1^ producing brighter marks than 50 mg L^-1^, suggests that at low winter growth rates, higher calcein concentrations are necessary to produce reliably detectable marks. Importantly, neither concentration caused mortality or observable behavioral effects, consistent with prior assessments of calcein as a benign fluorochrome in mollusks (e.g., Moran, 2000; Spires and North, 2022; van der Geest et al., 2011). The 100% survival rate and continued feeding activity during recovery indicate that the protocol can be applied without compromising animal welfare or confounding subsequent growth measurements.

Although shell marks are the primary target for growth studies, calcein fluorescence was also visible within the soft tissue of marked individuals and could serve as additional confirmation of exposure, though the persistence and utility of soft-tissue marks were not evaluated here and warrant further investigation.

Based on these results, we recommend a concentration of 100 mg L^-1^ with a 24-h immersion for marking *L. littorea* with calcein. However, marking success during winter will be strongly biased toward smaller, faster-growing individuals. Moreover, faint calcein fluorescence was detectable under a dissecting microscope in snails scored as unmarked with the field protocol, suggesting trace incorporation occurs even when growth rates are insufficient to produce field-detectable marks. For studies requiring marks across a broader size range, exposure during warmer months when growth rates are higher would improve marking success. Researchers employing this protocol in ocean acidification experiments should also consider that reduced calcification under low-pH conditions may suppress marking success through the same growth-rate mechanism documented here. Future work should evaluate seasonal effects on marking efficacy and determine minimum effective exposure durations, as shorter immersion times would reduce logistical constraints in both laboratory and field applications. Trials at higher concentrations could also be informative, given that 100 mg L^-1^ produced significantly brighter marks than 50 mg L^-1^ and caused no detectable adverse effects.

## Acknowledgements

Thank you to the staff at the Bowdoin College Schiller Coastal Studies Center and Biology Department for your help with laboratory assistance and logistical support. Special appreciation for the efforts of Heidi Franklin, Lisa Ledwidge, Holly Parker, Jaret Reblin, and Rachel Reuling.

